## Supporting Information for "Transient Formation of Supramolecular Complexes Between Hyaluronan and Oligopeptides at Submicromolar Concentration"

### NMR Meassurements

#### NOESY spectra on 8–15 KDa HA–nonapeptide mixtures

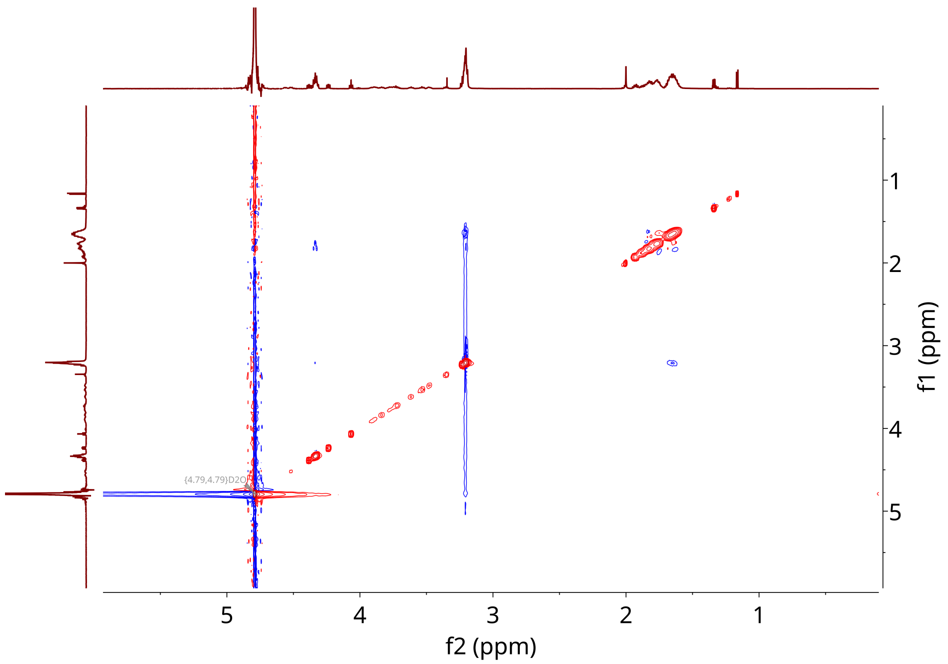


**Figure S1:** The NOESY spectrum for the HA–R9 mixture with 8–15 KDa HA.

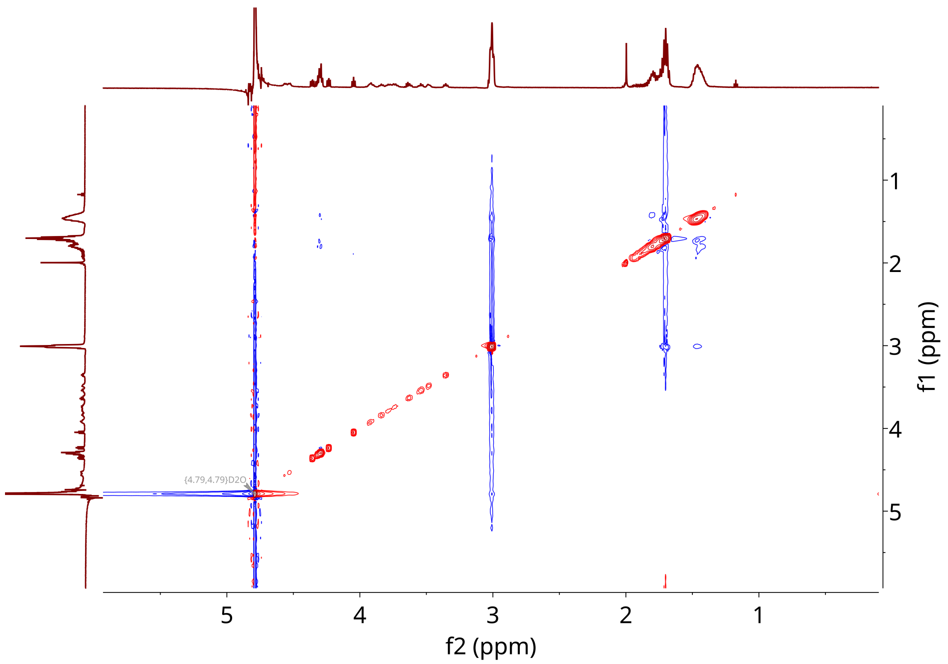


**Figure S2:** The NOESY spectrum for HA–K9 mixture with 8–15 KDa HA.

#### NMR spectra on supernatants from 1,375 KDa HA–nonapeptide mixtures

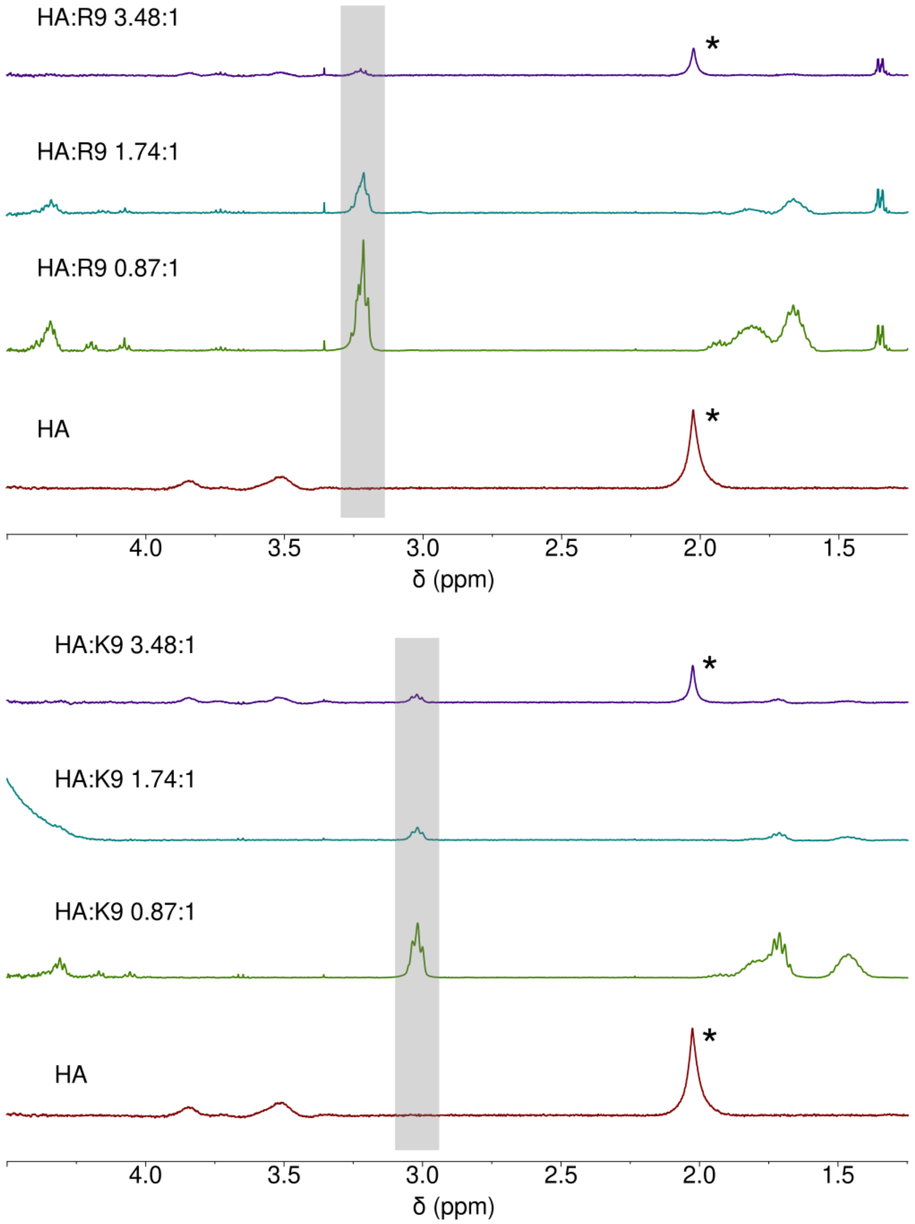


**Figure S3:** Stacked $^1$H NMR spectra of the supernatants at different HA dimer-to-amino acid ratios. The bottom trace corresponds to pure HA, while the others represent mixtures with R9 or K9 at varying concentrations. The gray-shaded region highlights the approximate area used for the integration of the peptide signal, selected for its clear and isolated peptide signal. One easily distinguishable signal, assigned to the acetamide group, is marked with an asterisk and was used to monitor HA's disappearance and subsequent reappearance in the solution. The spectra are produced in Mnova.[S1]

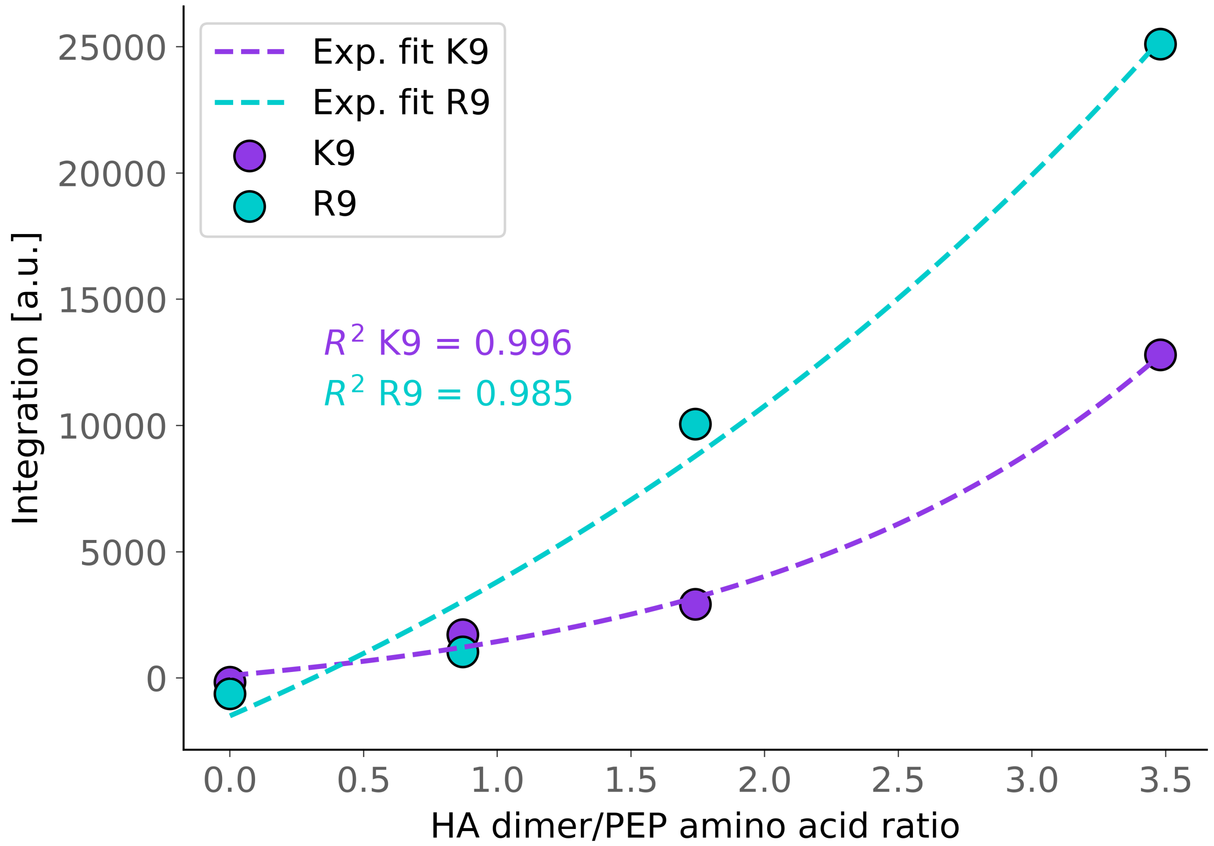


**Figure S4:** Integrals of the observable peptide signals in the supernatant (integration between 3.3–3.1 ppm for R9 and 3.1–2.9 ppm for K9, respectively) as a function of the HA dimer-to-amino acid ratio. The integrals were calculated in Mnova,[S1] with baselines being adjusted using Whittaker smoothing, which rendered clean, flat baselines in the integration region. The resulting data points where then fitted to an exponential function of the form $A \times e^{b \times x} + c$.

#### Supplementary solution images

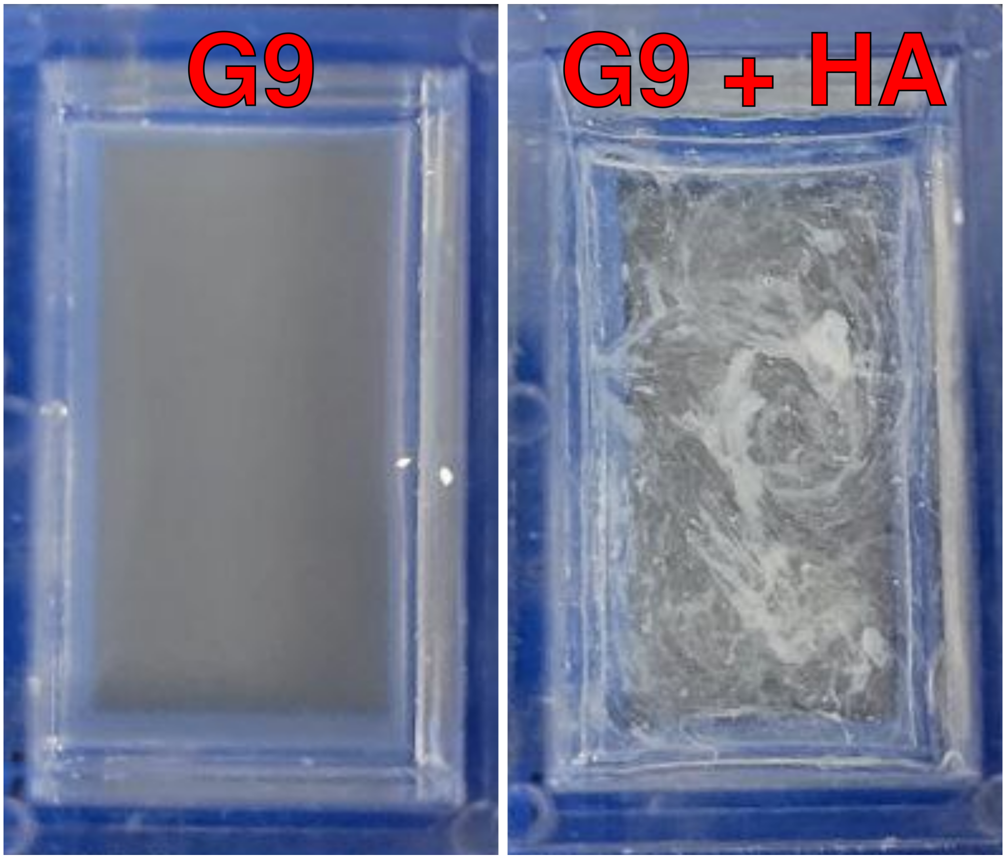


**Figure S5:** Unmagnified views of the G9 solution (left) and the G9 + 1,375 kDa HA mixture (right).

#### Supplementary AR-SHS data

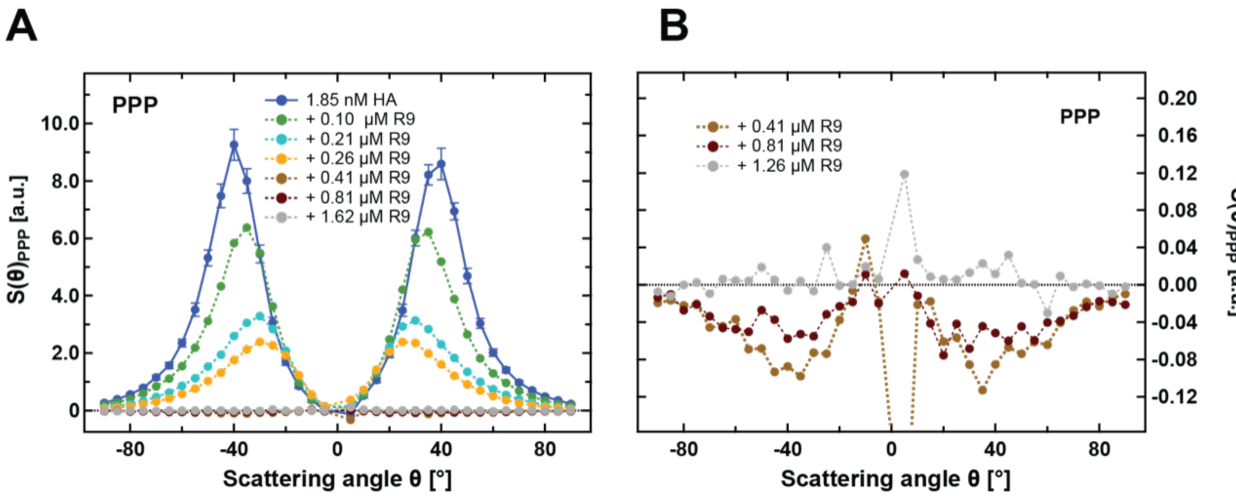


**Figure S6:** A) An extended series of normalized AR-SHS patterns in the PPP polarization combination of a HA solution (1,375 kDa) and HA–R9 mixtures. The HA solution has a concentration of 1.85 nM, and the mixtures include R9 at varying concentrations. The normalization is done following Eq. 1, using water as a reference for the HA solution and aqueous solutions of peptides as a reference for the mixtures. The concentration of peptides in the reference solution is equal to the one in the mixture. The effect of adding R9 to HA solutions on the pattern shape and intensity can already be seen at very low concentrations (0.05 μM). The effect is stronger with increasing R9 concentration. B) A zoomed-in view of the negative portion of the measured AR-SHS patterns at the highest R9 concentrations. A negative AR-SHS signal suggests at first that at the indicated concentrations, the mixtures induce less water orientation than the corresponding reference, here pure peptide solutions. Further studies would be necessary to fully understand the origin and the extent of this effect.

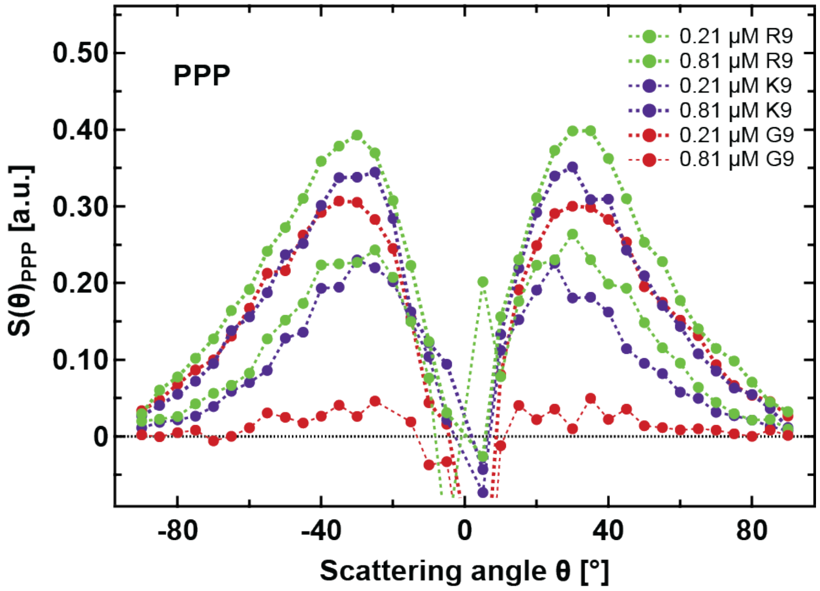


**Figure S7:** Normalized AR-SHS patterns in the PPP polarization combination of pure peptide solutions. The normalization is done following Eq. 1, using water as a reference. The concentration of peptides in the reference solution is equal to the one in the mixture.

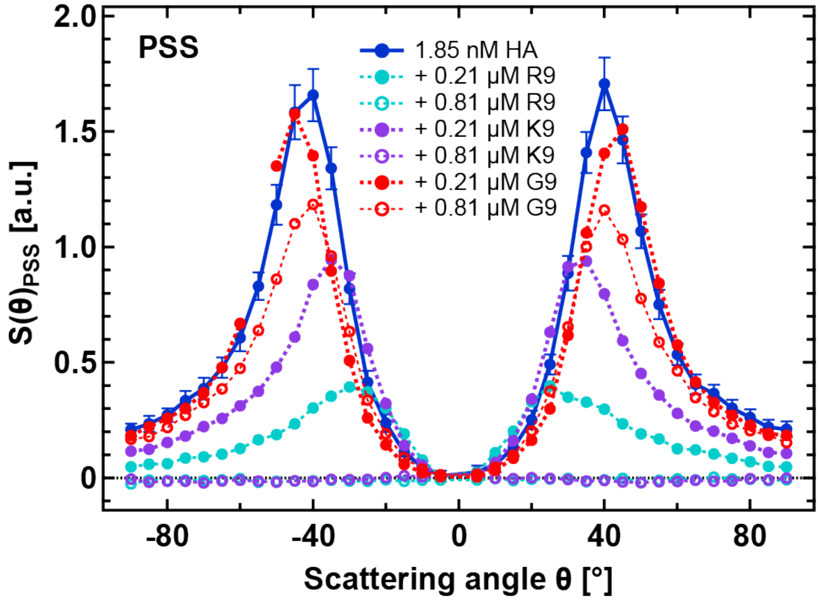


**Figure S8:** Normalized AR-SHS patterns in the PSS polarization combination of a HA solution (1,375 kDa) and HA–peptide mixtures. The HA solution had a concentration of 1.85 nM, and each peptide—either R9, K9, or G9—is added at two different concentrations: 0.21 µM and 0.81 µM. The normalization is done following Eq. 1, using water as a reference for the HA solution and aqueous solutions of peptides as a reference for the mixtures. The concentration of peptides in the reference solution is equal to the one in the mixture.

#### Conductivity and pH of AR-SHS solutions

The conductivity of each sample was measured and is reported in Table S1. Because we do not know the exact nature of the possible impurities, the measured conductivity cannot be converted to a known ionic concentration. However, one may estimate an upper limit for the possible ionic concentration by converting the maximum measured conductivity (5 µS/cm) using the molar ionic conductivity of an electrolyte with a typically low molar ionic conductivity, *e.g.*, KCl. This rough estimation indicates an upper limit of 20 µM of monovalent ions for the HA–peptide mixtures. Despite corresponding to a low amount of charged impurities, our optical experiment is extremely sensitive to extremely small amounts of charged impurities that could participate in the decrease in the AR-SHS signal intensity.

**Table S1:** Conductivity $\kappa$ and pH of studied HA and HA–peptide solutions for AR-SHS experiments.

| Sample | $\kappa$ [µS/cm] | pH |
|---:|---:|---:|
| HA 1.85 nM | 1.7 | 6.3 |
| HA 1.85 nM + R9 0.21 µM | 1.8 | 6.1 |
| HA 1.85 nM + R9 0.81 µM | 4.2 | 6.6 |
| HA 1.85 nM + K9 0.21 µM | 3.2 | 5.9 |
| HA 1.85 nM + K9 0.81 µM | 2.7 | 5.9 |
| HA 1.85 nM + G9 0.21 µM | 1.4 | 6.7 |
| HA 1.85 nM + G9 0.81 µM | 1.8 | 6.2 |

### Supplementary Molecular Dynamics Data

#### Solvent-accessible surface area

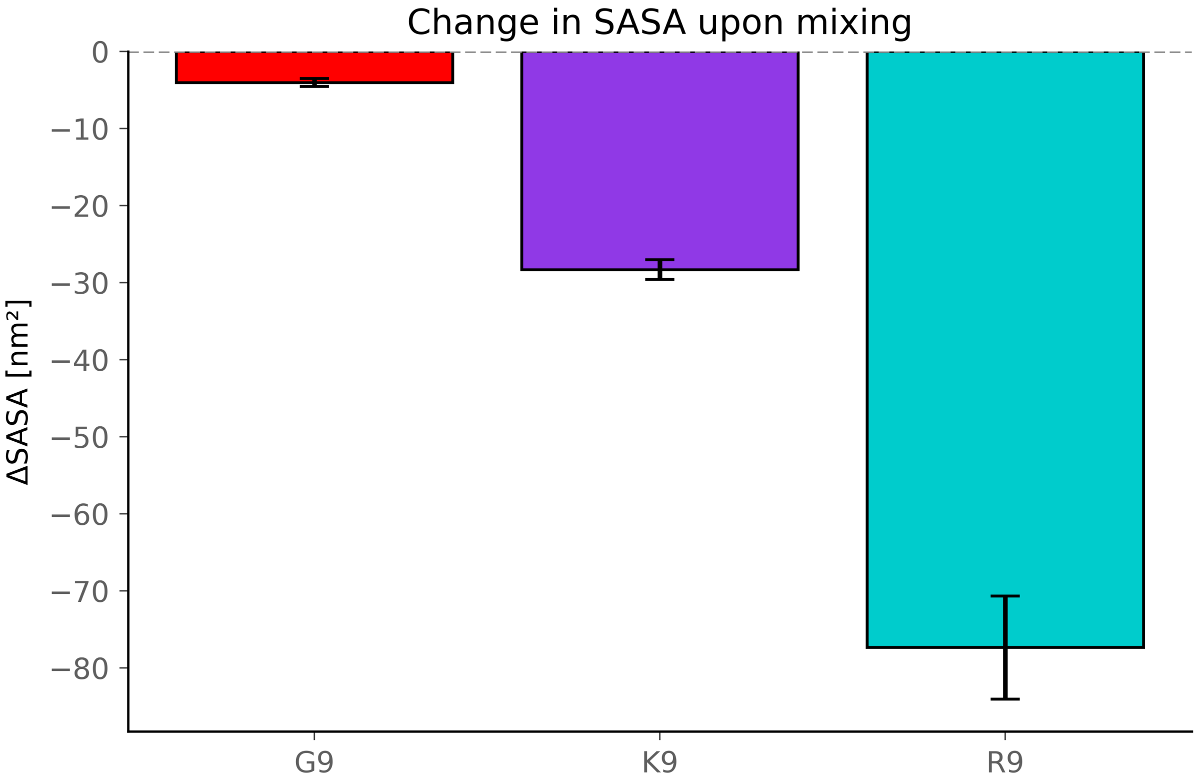


**Figure S9:** Change in the solvent-accessible surface area (SASA) of the solutes (HA and/or peptides) upon mixing, compared to the combined SASA from separate simulations of each component. The difference is calculated as $\Delta SASA = SASA_{mix} - (SASA_{HA} + SASA_{pep})$. The SASA values were calculated using `gmx sasa` tool in GROMACS software, version 2023.1,[S2,S3] with default parameters. The plot displays the average values across all replicas and the corresponding standard deviation.

#### Aggregate characterization

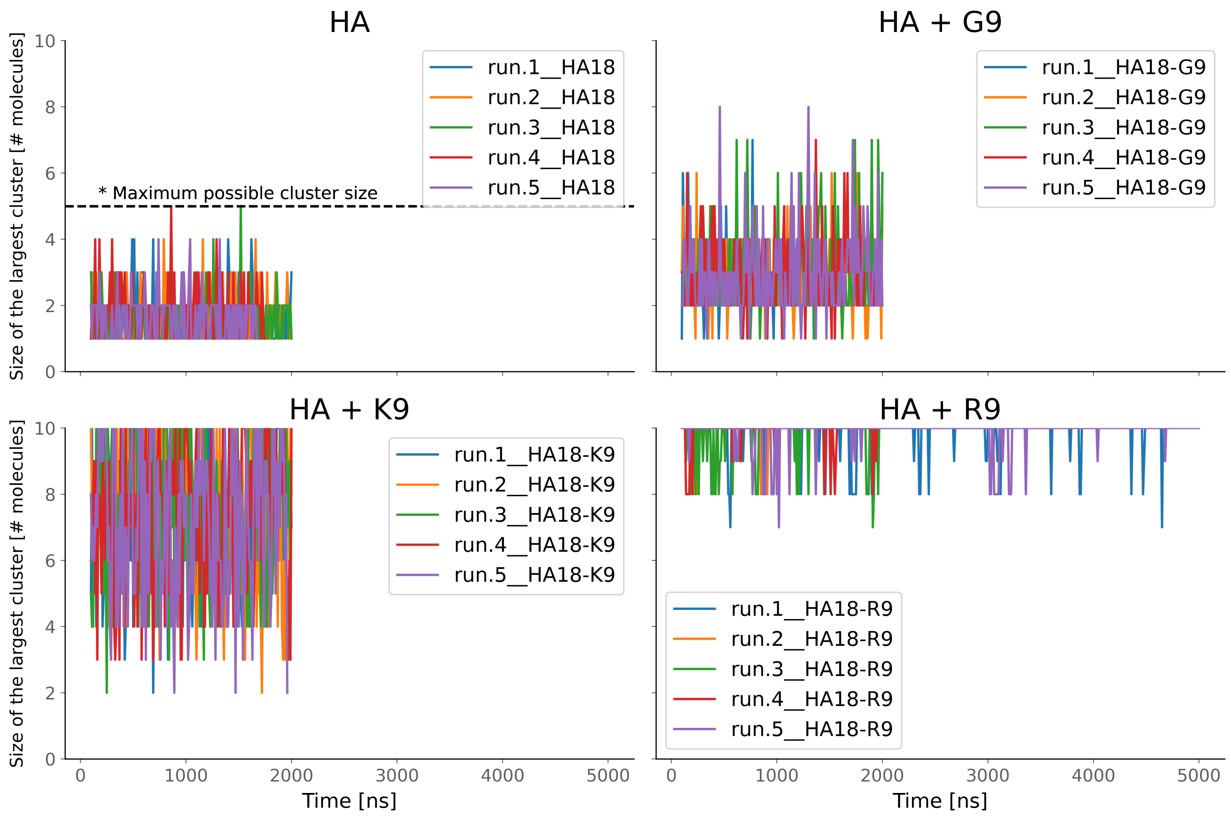


**Figure S10:** Time evolution of the size of the largest cluster. All replicas for each system type are shown. The size of the largest cluster is defined as the number of solute molecules (HA and/or peptides) in continuous contact, with a cutoff of 3.5 Å.

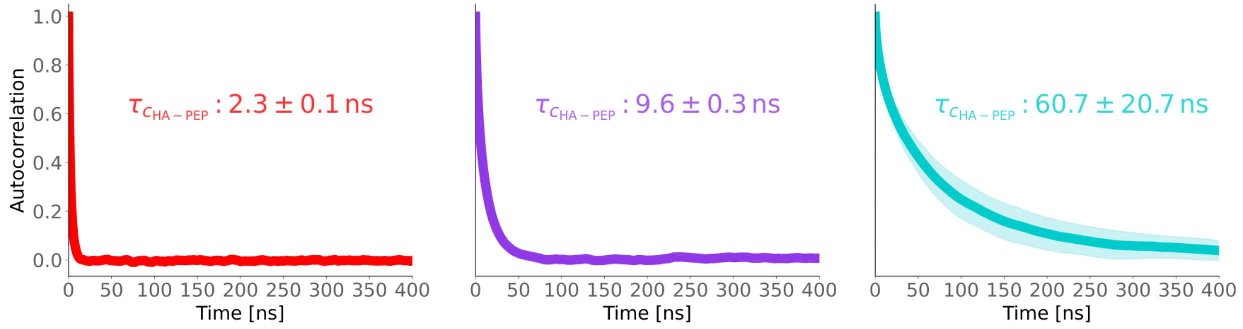


**Figure S11:** HA–peptide distance autocorrelation function. The autocorrelation characteristic times are calculated by fitting the data to an exponential function of the form $e^{-t/\tau_c}$, where $\tau$ is the lag time. The standard deviation—calculated for all simulation replicas and shown as a shaded area—is notable only for HA–R9 systems.

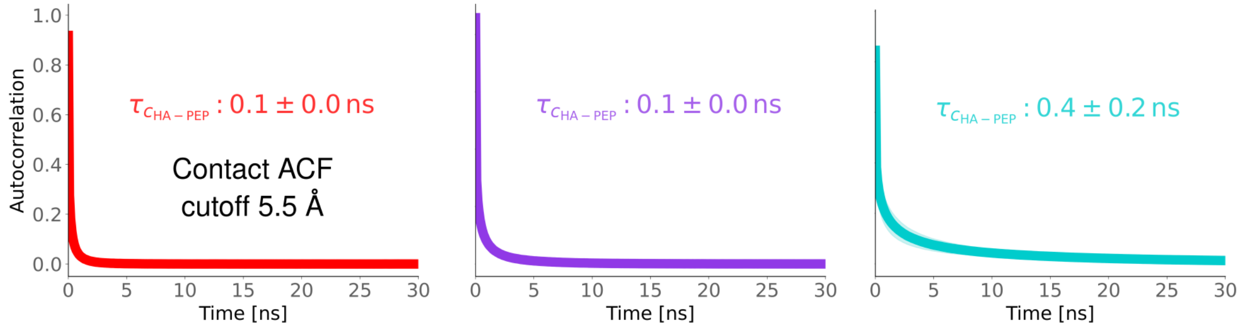


**Figure S12:** HA–peptide contact autocorrelation function. The HA–peptide residues center of mass distances are converted to contacts with a cutoff of 5.5 Å. The autocorrelation characteristic times are calculated by fitting the data to an exponential function of the form $e^{-t/\tau_c}$, where $\tau$ is the lag time. The standard deviation is calculated for all simulation replicas and shown as a shaded area.

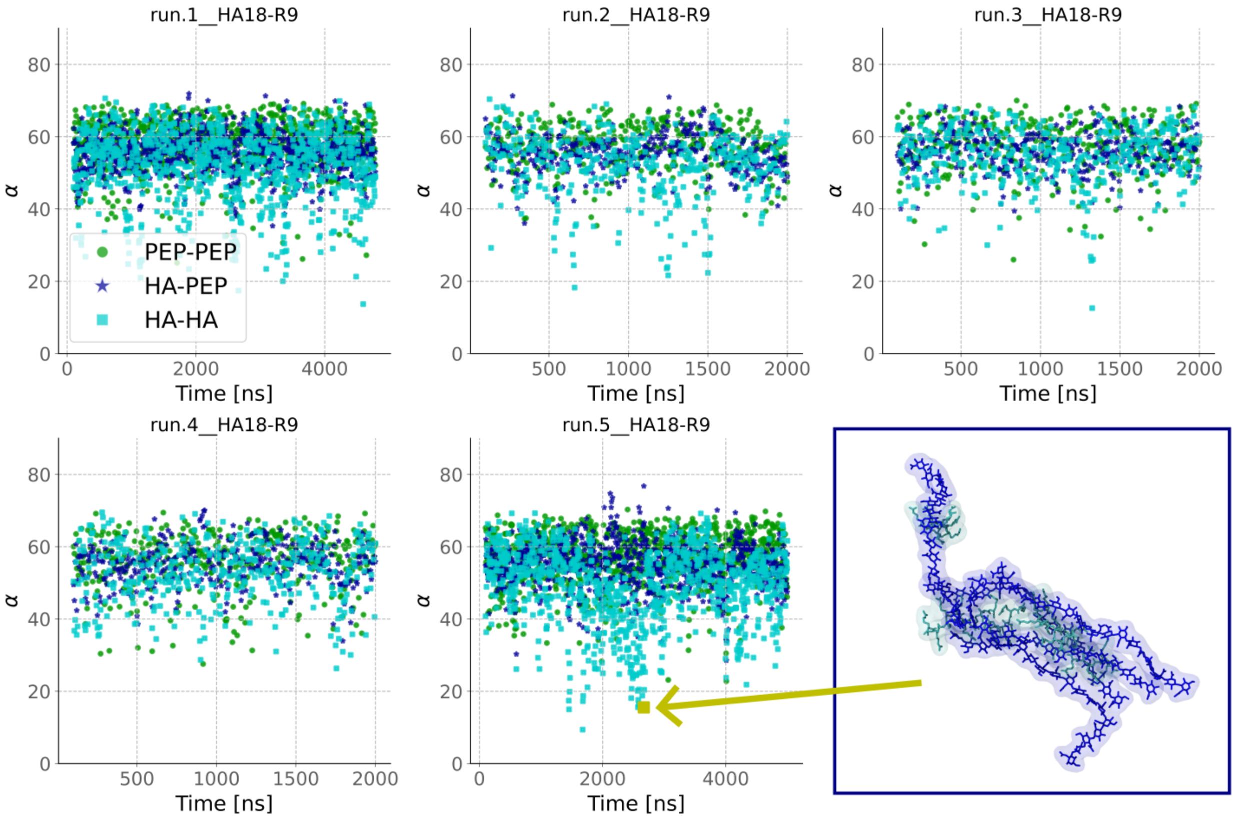


**Figure S13:** Time evolution of the average angles between the molecules in the HA–R9 systems. All five replicas are shown independently, and the HA–HA, HA–PEP, and PEP–PEP are shown. The angles shown are the average angle between all pairs of molecules of a given type. The molecular axis used for the angle analysis was determined using singular value decomposition. A snapshot from a frame with low HA–HA angle values is included to illustrate the bundled structures associated with these low-angle configurations. These bundles form and dissociate spontaneously throughout the simulations and are not observed in the K9 or G9 systems. The HA molecules are shown in blue, while the R9 is shown in cyan. A yellow dot and arrow highlight the specific frame from which the snapshot was taken.

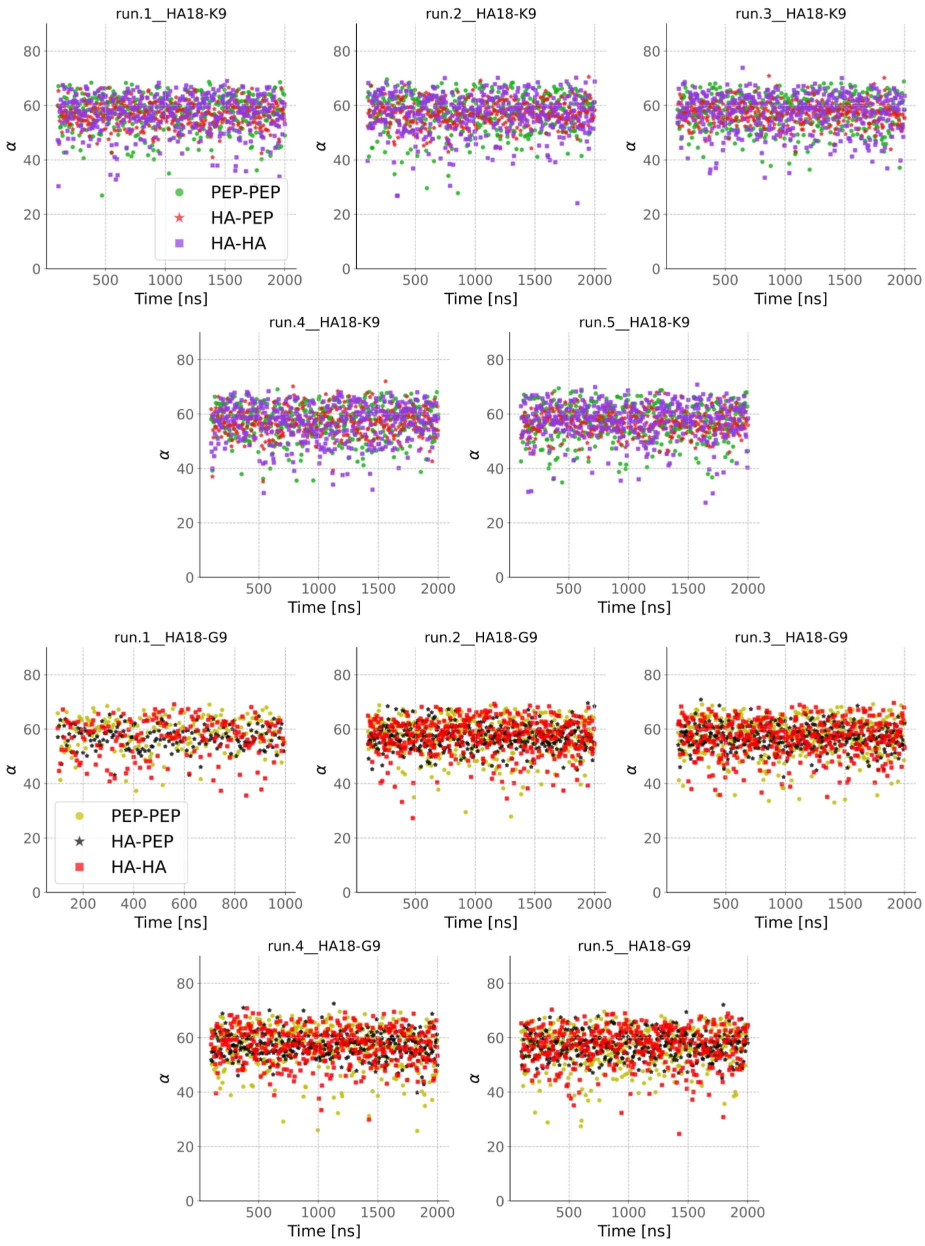


**Figure S14:** Time evolution of the average angles between the molecules in the HA–K9 (purple) and HA–G9 (red) systems. All five replicas are shown independently, and the HA–HA, HA–PEP and PEP–PEP are shown. The angles shown are the average angle between all pairs of molecules of a given type. The molecular axis used for the angle analysis was determined using singular value decomposition.

#### Orientation of water molecules with respect to the solutes

Water orientation is calculated as the angle between the water bisector vector and the vector pointing from the water oxygen to the nearest solute atom. Since water is in the liquid state, it is highly mobile, and when averaging over many configurations, an angle of $90°$ indicates random orientation. Angles less than $90°$ indicate that, on average, the water hydrogen atoms are oriented toward the solute. In comparison, angles greater than $90°$ suggest that the water oxygen tends to be closer to the solute.

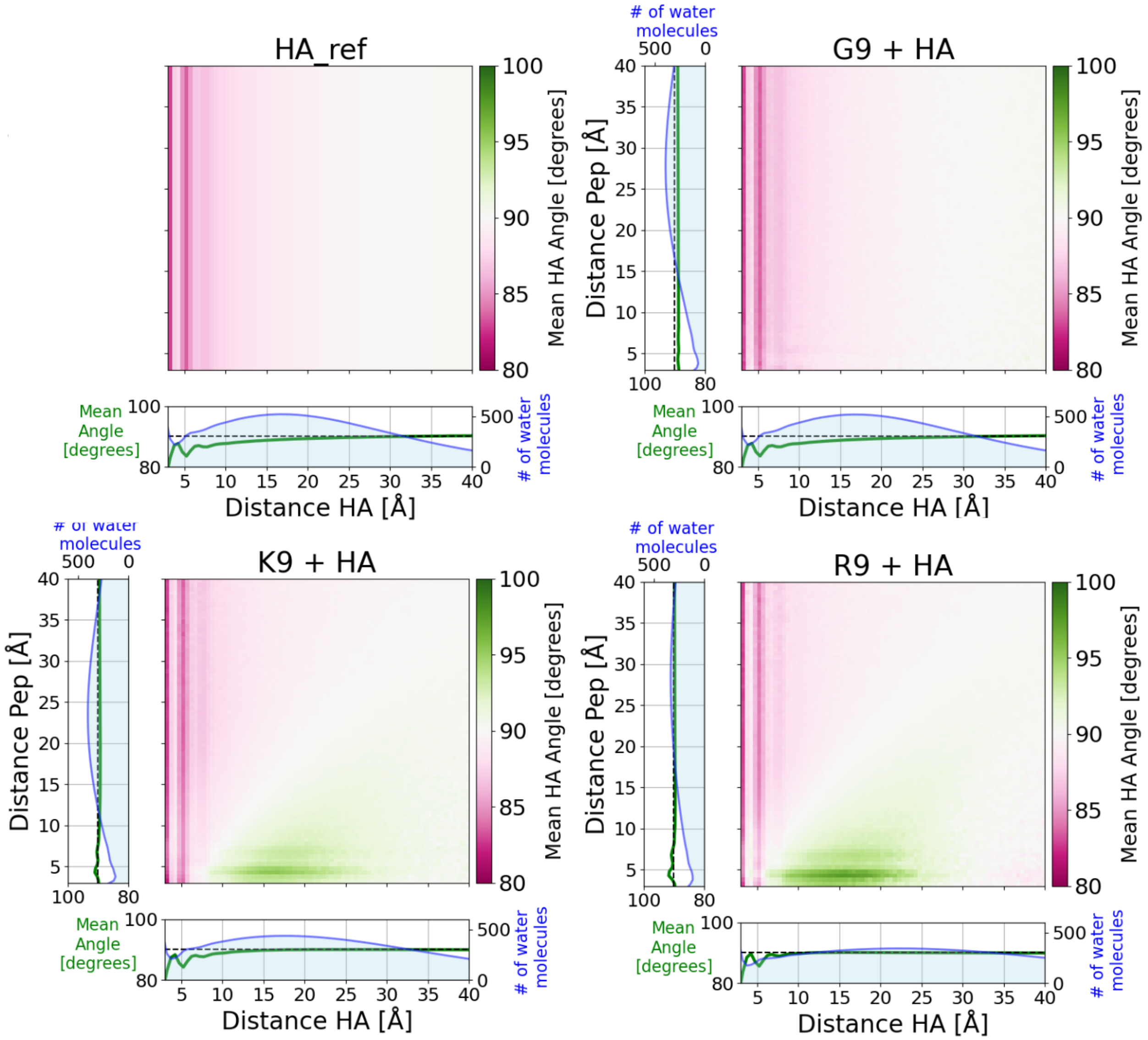


**Figure S15:** Absolute average water orientations with respect to HA as a function of the distance to both HA and peptides. The orientation of water molecules was calculated as the angle between the water bisector and the vector connecting the water oxygen to the nearest atom of an HA molecule. These data are used as a reference for the normalization in Figure 4B in the main text. In the pure HA system (top left), there are no peptides, so the vertical axis is constant. Marginal plots show the average angle in each bin and the average number of water molecules per frame in that bin.

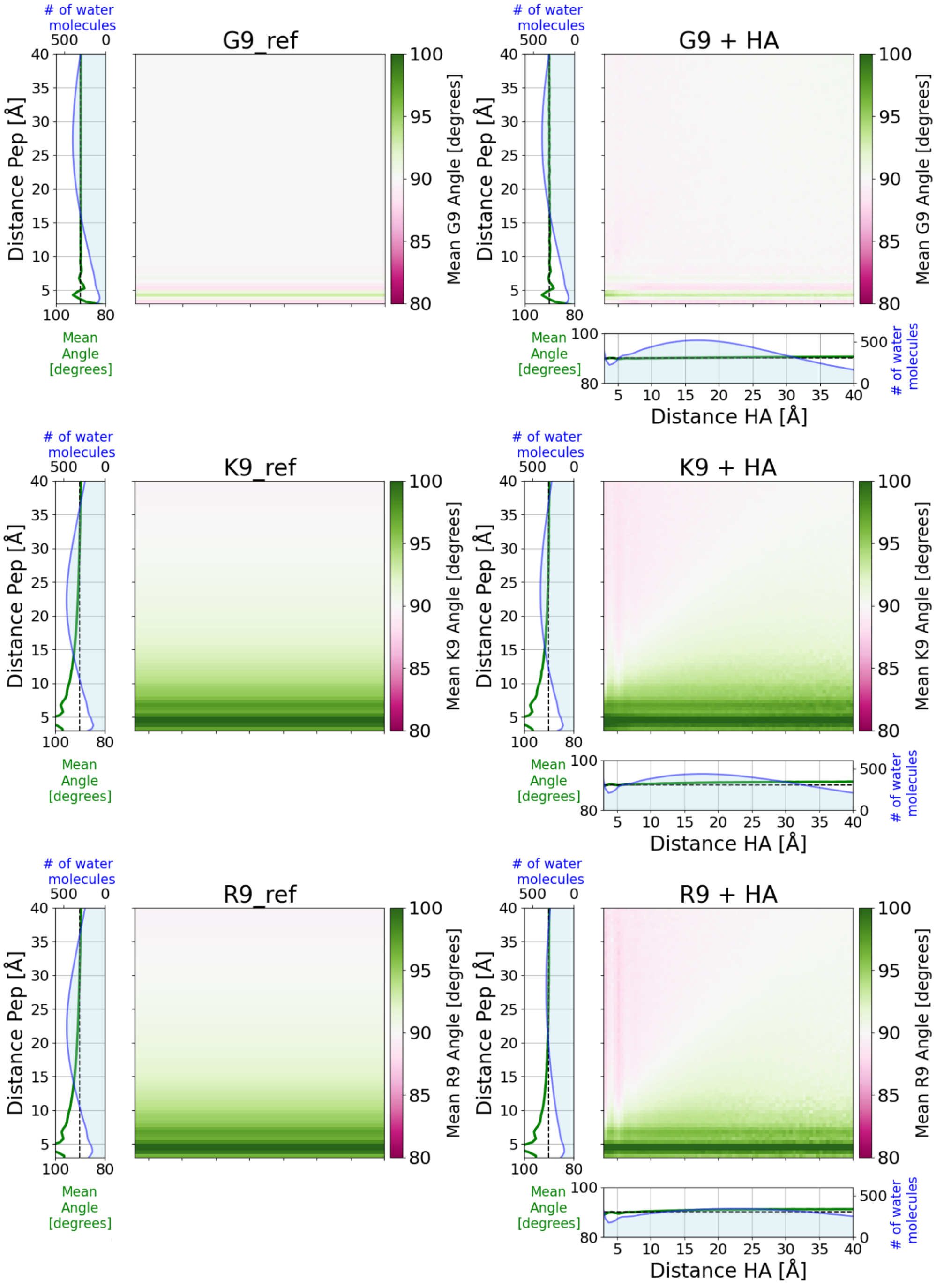


**Figure S16:** Absolute average water orientations with respect to peptides as a function of the distance to both HA and peptides. The orientation of water molecules was calculated as the angle between the water bisector and the vector connecting the water oxygen to the nearest atom of a peptide molecule. These data are used as a reference for the normalization in Figure 4C in the main text. In the pure peptide systems (left column), there is no HA, so the horizontal axis is constant. Marginal plots show the average angle in each bin and the average number of water molecules per frame in that bin.

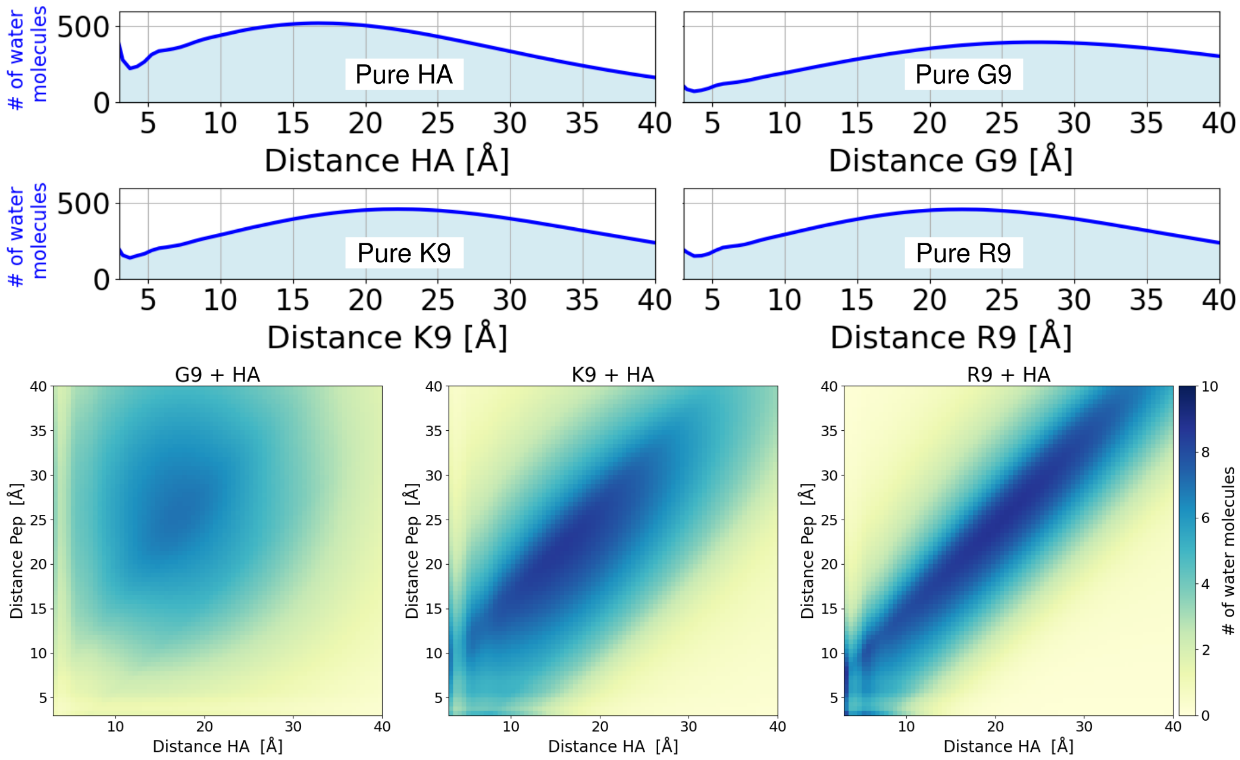


**Figure S17:** Number of water molecules as a function of distance to HA or the corresponding peptide. The two upper rows show the distribution of water molecules in pure solutions. The 2D plots show the same for the HA–peptide mixtures.
